# Heat stress sensitizes zebrafish embryos to neurological and cardiac toxicity

**DOI:** 10.1101/2024.09.03.609146

**Authors:** Anna-Mari Haapanen-Saaristo, Noora Virtanen, Elena Tcarenkova, Katri Vaparanta, Minna Ampuja, Eeva-Riikka Vehniäinen, Ilkka Paatero

## Abstract

Global warming increases the risk of dangerous heat waves, which may have deleterious effects on humans and wildlife. Here, we have utilized zebrafish embryos as a model to analyse heat stress and effect of chemical compounds on responses to heat stress. The temperature adaptation limit of zebrafish embryos was 37°C in behavioural test and 38°C in cardiac test. Polyaromatic hydrocarbon phenanthrene completely blocked the behavioural adaptation to heat stress. Interestingly, the cardiotoxic effects of lapatinib, phenanthrene and paclitaxel were induced by heat stress. Taken together, our data indicates that motility and cardiac function of zebrafish embryos can be utilized as a model to analyze modulatory effects of compounds on heat stress.

**Highlights:**

- Zebrafish embryos can be utilized as an in vivo model for acute heat stress
- Phenanthrene inhibited motility increase upon heat stress
- Cardiotoxicity of lapatinib, paclitaxel and phenanthrene was potentiated by heat stress

## Introduction

Heat waves cause excess heat exposure and can be highly lethal. Heat stress increases the metabolic rate, and thus increases the demand for oxygen and increased cardiac output is needed for increased transfer of oxygen to tissues. The heart can adapt to high temperatures to a certain extent, but extreme heat exposure results in cardiovascular disease such as heart failure, cardiac arrest and arrythmias [1] and cardiovascular disease accounts for up to 90% of heat wave mortality [2]. Excess exposure to heat results in hyperthermia or heat stroke. In humans, the mortality of heat stroke can exceed 50% and there are no specific pharmacological treatments for heat stroke the most common therapeutic action being cooling [3].

Humans and other mammals adapt and regulate their body temperature, but excessive heat exposure results in life-threatening heat stroke where body temperature regulation cannot cope with excess heat resulting in increased body temperature [3]. Heat exposure is not problem only to humans but heat waves have also severe effects in wildlife [4]. Cold blooded (poikilothermic and ectothermic) animals such as fish cannot adapt to the heat exposure with body temperature regulation but are directly affected by increased ambient temperatures. Due to global climate change, the heat exposure is increasingly significant factor both in humans and animals [5,6].

Experimental elucidation of factors affecting heat stroke and associated heart failure requires efficient research models. Zebrafish models have been turned out to be efficient and reliable research models for heart failure and cardiovascular biology [7]. As zebrafish embryo heart clearly responds to increased temperature [8], we tested whether it could be used to model also heat induced heart failure and to analyse potential of heat stress in sensitization to cardiac effects of chemical compounds with potential cardiotoxicity.

## Materials and methods

### Treatment with chemical compounds

All reagents were purchased from Sigma-Aldrich and dissolved in DMSO. Glucocorticoid receptor agonist dexamethasone, anti-cancer drugs doxorubicin, lapatinib and paclitaxel, and polycyclic aromatic hydrocarbon (PAH; an air pollutant) phenanthrene were used at 10 µM, and positive control beta-blocker propranolol at 1 µM concentration. As a negative control, 1% DMSO (maximum solvent concentration in the treatments) in E3 medium was utilized. The embryos were incubated with the compounds for 1 hour before analysis.

### Zebrafish husbandry and mating

Adult zebrafish were housed in Aqua-Schwarz PP-module stand-alone system under license MMM/465/712-93 (Ministry of Agriculture and Forestry of Finland) at 12h:12h light-dark cycle. The embryos were obtained by natural spawning in 1.7L sloped breeding tanks (Tecniplast, IT). After spawning, the embryos were collected and transferred into E3 medium in petri dishes and incubated at 28.5°C until used in experiments.

### Motility analyses

At 4 days post-fertilization (dpf), the zebrafish embryos were exposed to chemical compounds in E3 medium. The tested drugs were dissolved in DMSO and applied into wells of multiwell plate each containing one embryo. The motility assays were carried out using DanioVision (Noldus IT) instrument equipped with temperature control unit and Ethovision XT 14 software. In the temperature finding experiment, the temperature was first stabilized into 28.5°C, and then ramped up to 40°C while simultaneously tracking movement and temperature. Sample temperatures were measured and logged with HOBO MX2201 Pendant water temperature data logger (Onset Computer Corporation, Bourne, MA, USA). In the drug exposure experiments, the embryos were analysed in 28.5°C and 38°C. The movement was tracked for an hour.

### Brightfield cardiac imaging

To analyze cardiac function, the zebrafish embryos at 4 dpf were exposed to increasing temperature (from 28.5°C to 40°C) and imaged. To analyze the cardiac function at physiological temperature and heat stress, the treated embryos were exposed to 28.5°C or 38°C for 5 minutes and imaged. OxyGenie instrument (Baker Ruskinn) was utilized for temperature control. The embryos were anesthetized with 200 mg/l MS-222 before heat exposure and imaging. AxioZoom V.16 (Zeiss) stereomicroscope with 1.0x Plan ApoZ (NA.0125) objective and Hamamatsu sCMOS Orca Flash4.0 LT + camera with fast time-lapse imaging at 51 frames per second (fps) were utilized. Movies were converted to .avi format and analyzed using DanioScope (Noldus IT) software.

### High-resolution intravital imaging

For high resolution imaging we used multi-point confocal microscopy (3i CSU-W1 spinning disk, 3i, Denver, Colorado). Objective LD Plan-Neofluar 40x/0.6 Corr Ph2 M27 with working distance of 2.9 mm at 0.75 glass was used. PRIME BSI High Resolution BSI Scientific CMOS and Hamamatsu sCMOS Orca Flash4.0 cameras were used. Both systems provided high sampling capacity with 6.5×6.5 µm pixel size. Excitation with 488nm and 561nm laser and quad band (440/40nm, 521/21nm, 607/34nm, 700/45) emission filter were used. A low exposure time (30-50 ms) was used to achieve the most real-time imaging possible. Incubation chamber (OKOlabs, Napoli, Italy) was used during the imaging to maintain the temperature at desired level. Embryos of strain Tg(cmlc2:EGFP, kdrl:mCherry) were mounted on glass bottom dish (Cellvis #D35-10-1.5-N), anesthetized with tricaine (MS222, Sigma Aldrich, Germany) and mounted in agarose (Sigma Aldrich, Germany) to prevent any extra movements. To further analyze cardiac function, the immobilized embryos were exposed to a temperature that steadily increased from 28°C to 37°C over 10 minutes. Imaging was performed for each embryo at both temperature points to assess the effects of this gradual heating on cardiac performance. Diameter of ventricle was measured along short axis in both diastole and systole, and fractional shortening (FS) calculated as FS(%) = 100% * (Diastole - Systole)/Diastole.

### Statistical analyses

Statistical analyses were carried out using GraphPad Prism. The normality and homoscedasticity assumptions were tested with Anderson-Darling, D’Agostino-Pearson omnibus, Shapiro-Wilk, Kolmogorov-Smirnov, Brown-Forsythe and Bartlett’s tests and parametric or non-parametric testing was chosen accordingly. For non-parametric testing, Kruskal-Wallis ANOVA and Dunn’s multiple comparison test was utilized. For parametric testing, one-way ANOVA and Dunnett’s multiple comparison test was utilized. For parametric testing of fractional shortening, a paired t-test was carried out. The method of ROUT at alpha level 0.05 was utilized for outlier removal. The line in the boxplot corresponds to the median value, the box to the interquartile range and whiskers to the whole range of values.

## Results

The organisms can physiologically adapt to range of ambient temperatures and to model heat stress in zebrafish embryos we first aimed to identify higher end of temperature range where the animals were still able to adapt their physiological performance. To analyze temperature effects and fast adaptation capabilities of zebrafish embryos, we carried out behavioural analysis using fast measurements of motility and temperature (Fig. 1A), as the motility is often used as a general measure of physiological performance of the animal [9]. The capability of zebrafish embryos to adapt was diminished around 37°C (Fig. 1A), followed by slow decline in motility. To more formally define the adaptation temperature limit, we calculated first derivatives of swim distance and velocity, and determined the point where 1^st^ derivative dropped below 0. With this approach, the adaptation limit was at 36.8°C (Fig. 1B).

**Figure 1.**
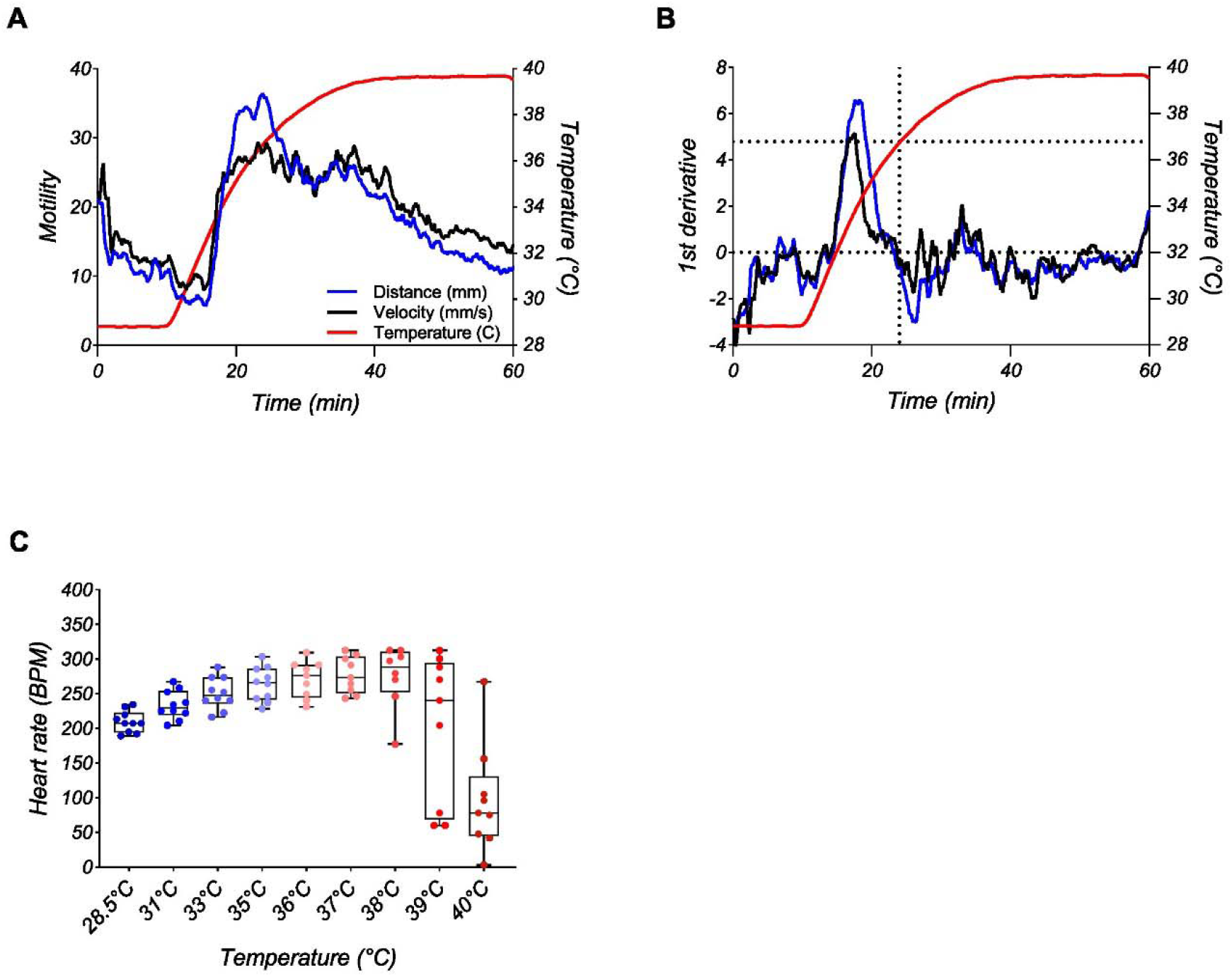
Optimization of temperature for heat stress experiments. A and B) Motility of zebrafish embryos during rising temperatures. A) Distance and velocity curves smoothened (moving average of 1-minute time window). B) 1^st^ derivative of motility curves. Mean value of 23 embryos were plotted, error bars were omitted due to clarity. C) Heart –rate measurements of embryos at different temperatures. Mean and standard error of 8-10 embryos / temperature are plotted, individual data points are shown in the graph.

Since zebrafish heart rate is known to be affected by the temperature [8], we next confirmed the previously determined temperature limit of adaptation by measuring heart rate of embryos at different temperatures. Expectedly, the heart rate increased with temperature until reaching maximum plateau value at 38°C (Fig. 1C). In temperatures above 38°C, the heart rate dropped (Fig. 1C) indicating severe cardiac heat stress and loss of physiological adaptation capability. Taken together this data indicates that temperature limit for zebrafish embryos to adapt to short-term heat stress was around 37-38°C, depending on the used measurement.

This data suggested that heart function declines with acute exposure to high temperatures. To get more detailed insights into the adaptation capability of the heart, we carried out fast fluorescence time-lapse imaging of the beating ventricle of transgenic zebrafish embryos carrying *cmlc2:EGFP* transgene expressing enhanced green fluorescent protein (EGFP) in myocardium (Fig. 2A). Imaging at high temperature of 37°C (just below the adaptation limit) indicated that overall ventricular anatomy was normal and ventricle contraction was not qualitatively compromised. Measurement of fractional shortening in few samples did not indicate strong alteration of fractional shortening by short-term incubation at 37°C (Fig. 2B). This indicated that short-term cardiac response to heat stress occurred mainly at the level of heart rate.

**Figure 2.**
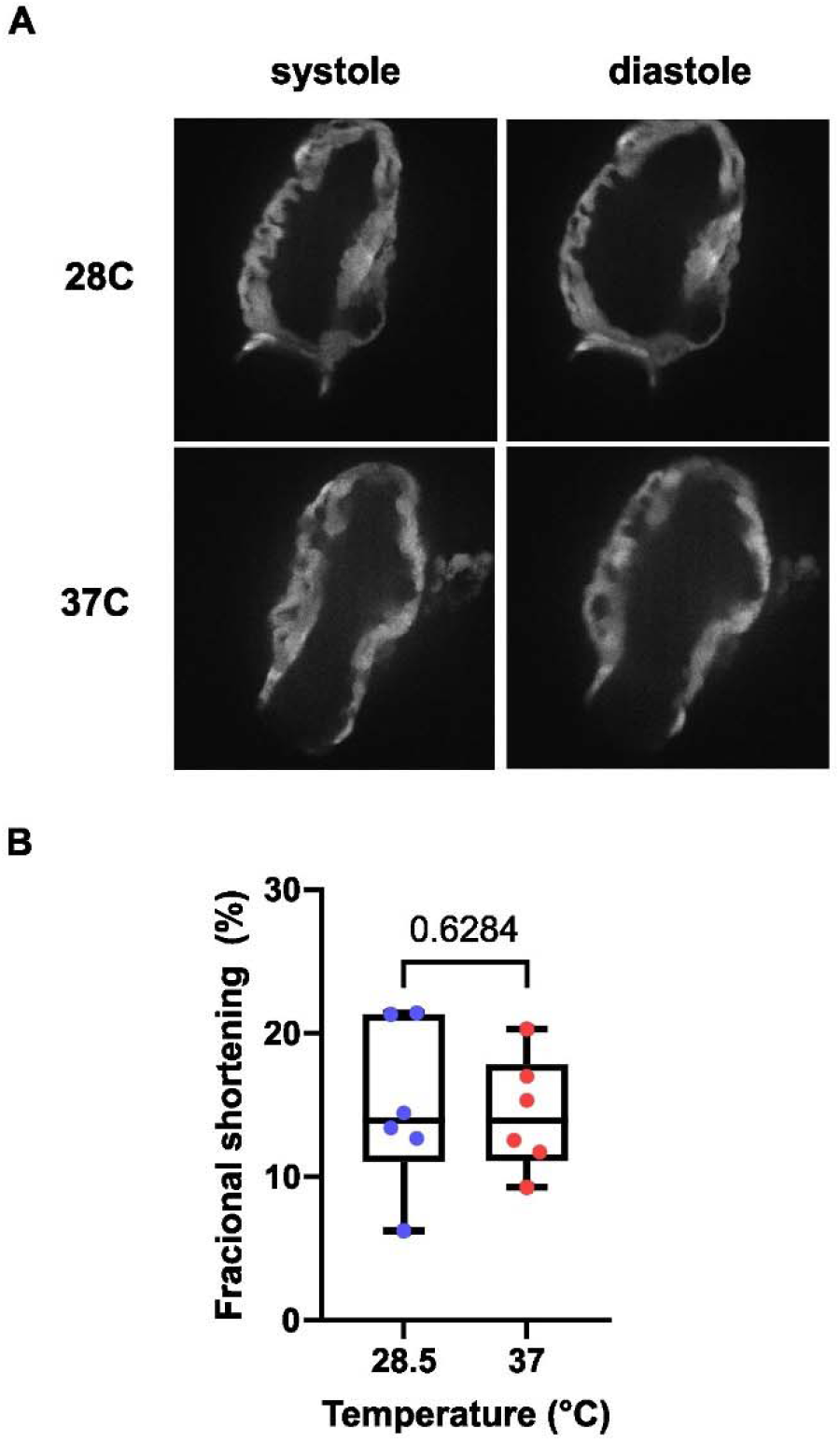
High-resolution imaging of heart-function. A) Spinning disk confocal imaging of EGFP fluorescence at 28.5C and 37C. The overall gross morphology of myocardium was normal. B) Quantification of ejection fraction from the high-speed fluorescence movies. One dot corresponds to one embryo, n=6.

We hypothesized, that heat stress would sensitize to effects of cardiotoxic chemicals. To analyze potential of acute heat stress to alter sensitivity we exposed zebrafish embryos to various compounds known to possess cardiovascular effects. First, we carried out motility analyses both in normal and heat stressed conditions to gain insights into overall and neuromuscular effects of the compounds in zebrafish embryos. In the normal temperature doxorubicin and lapatinib increased motility of the zebrafish embryos (Fig. 3A). When the embryos were subjected to heat stress, increased motility caused by doxorubicin and lapatinib persisted (Fig. 3B). In addition, zebrafish embryos subjected to phenanthrene decreased their motility in 38°C and dexamethasone treated embryos had tendency towards decreased motility (Fig. 3B). The distance moved is essentially a product of speed and duration. The effects on distance were mainly explained by altered duration of motility (Fig. 3C and D) indicating behavioural response to heat stress, and using this measure dexamethasone produced clearer effects. To analyze adaptation capability and to overcome changes in baseline activity, we calculated ratios of motility in 28.5°C and 38°C (Fig. 3E). The data indicated that in addition to changes in overall motility, also capability to respond to temperature increase was reduced in phenanthrene-treated embryos.

**Figure 3.**
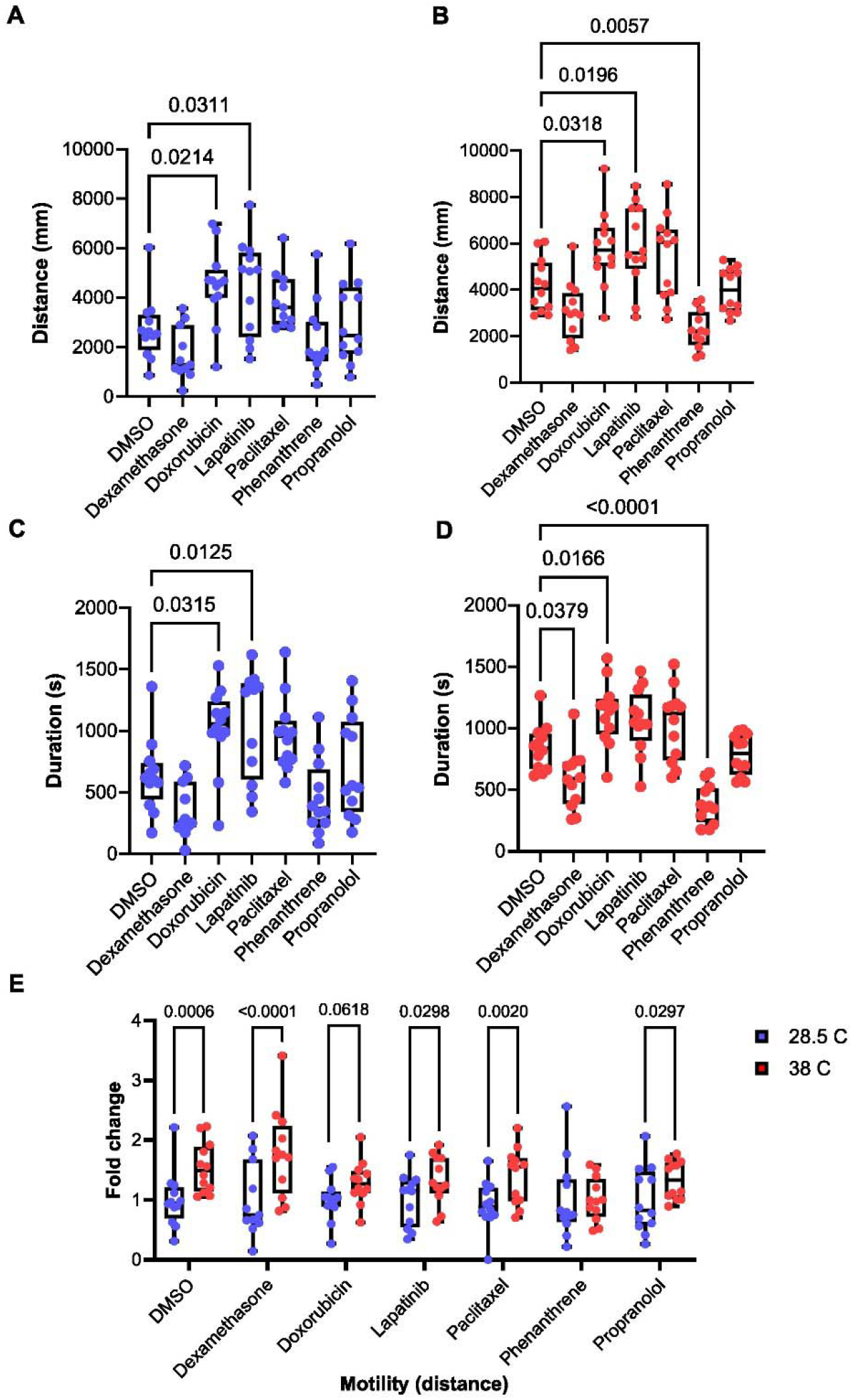
Analysis of short-term motility defects cause by inhibitors. Zebrafish embryos were treated with different compounds and subjected first to 28.5°C and then 38°C temperatures for motility analysis using Daniovision instrument. A) Distance of movement in 28.5°C. B) Distance of movement in 38°C. C) Duration of movement in 28.5°C. D) Duration of movement in 38°C. E) Adaptation capability was measured as ratio of distance moved in 38°C and 28.5°C. N=11 to 12 for each treatment.

Next, we looked at cardiac effects in acute exposure to tested compounds as the temperature has clear effect on cardiac physiology of zebrafish embryos. At normal temperature of 28.5°C only propranolol had effects on the heart rate (Fig. 4A). When the embryos were exposed to heat stress, also lapatinib, paclitaxel and phenanthrene had significant effects on the heart indicating vulnerability of heat stressed heart to these compounds (Fig. 4B). Taken together, this data indicated that heat stress may promote acute neuromuscular and cardiovascular toxicity.

**Figure 4.**
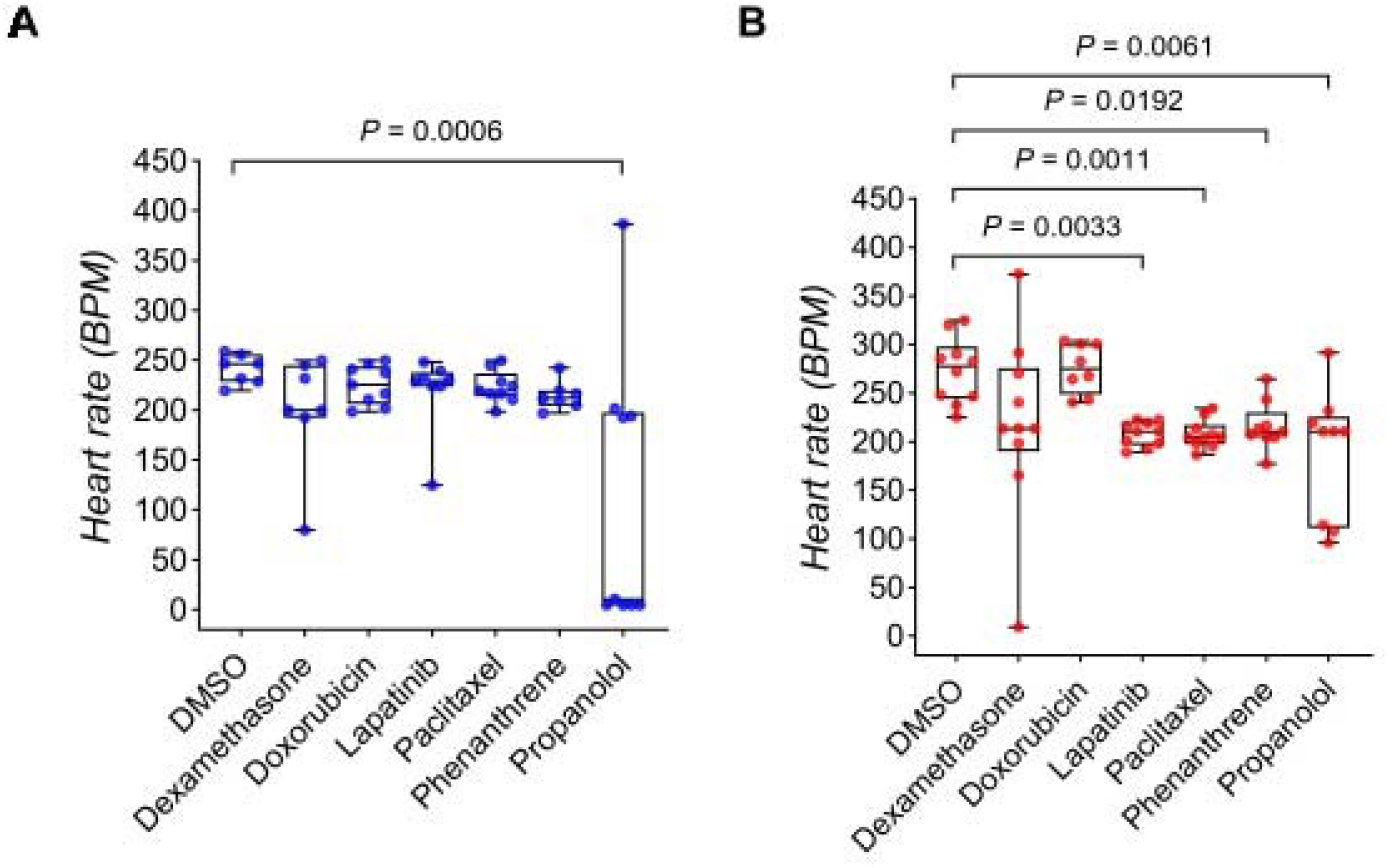
Acute cardiac defects of embryos exposed to chemical compounds and heat. Zebrafish embryos at 4 dpf were treated with the indicated compounds and exposed to either physiological temperature of 28.5°C (A) or at 38°C (B). The heart rate of the embryos was measured with fast time-lapse brightfield imaging. BPM: beats per minute. One dot corresponds to one embryo.

## Discussion

We observed that heat stress may alter the sensitivity of zebrafish embryos to various compounds. This data can be interpreted in two different views. It can be approached either from an ecotoxicological or human health point of view. As these views are quite different, we discuss these views here separately.

In our experiments, the zebrafish embryos responded to acute heat stress with increased locomotion and heart rate. This increase was observed until 37-38°C, with subsequent reduction in function. This indicates that short-term adaptation limit is on that region. The logic is somewhat analogous to critical thermal maxima (CT_max_) test used in studies with many fish species [10]. CT_max_ in adult zebrafish can be as high as 41°C [11], and the discordance between adult CT_max_ and our observations in embryos may be due to different methodology, or that zebrafish embryos are more sensitive to high temperatures.

Zebrafish embryos seem to respond to heat stress by increasing their heart rate rather than increasing the ejection fraction. Also in adult zebrafish, increased heart rate is the main factor behind increased cardiac output with rising temperature [12]. It is also possible, that longer exposures could lead to additional effects on morphological or physiological level which potentially could also translate into effects in different parameters in cardiac physiology, including ejection fraction.

Temperature often exacerbates the toxicity of chemicals in aquatic environments (reviewed in [13,14], and chemical exposure may lead to diminished ability of fish to cope with thermal stress [15]. This can be seen in our results as well, as one-hour exposure to lapatinib, paclitaxel, phenanthrene or propranolol inhibited the increase in heart rate caused by acute heat stress. Whereas longer exposures to phenanthrene and propranolol are known to cause bradycardia in zebrafish embryos at µM concentrations [16–18] there is no literature for cardiac effects of lapatinib and paclitaxel. We used short-treatments with high-doses, which may not be directly environmentally relevant exposure. However, the longer-term risks can be still estimated based on shorter-term exposures [19] and typically lower concentrations are more effective upon longer exposures.

Doxorubicin and lapatinib increased the motility of zebrafish embryos at both temperatures, suggesting they are anxiogenic or irritating. Phenanthrene has also earlier been shown to lower swimming distance in zebrafish embryos, but only after several days of exposure [20]. Dexamethasone had borderline effects and reduced swim duration statistically significantly but not swim distance upon heat stress. In adult zebrafish, 7-day treatments with dexamethasone have resulted in reduced swim performance [21]. It must be noted that these effects in our experiments were seen already after one hour of exposure. In addition, also impact of paclitaxel, lapatinib and phenanthrene on heart rate was observed in heat stress already after short exposure. It is plausible that with longer exposure, effects could be seen at lower concentrations. All in all, the results suggest that the interplay between environmental contaminants and heat stress may be detrimental to fish embryos. Although we have here analysed only motility and heart rate, it is likely that this interplay of chemicals and heat stress is present also in other physiological processes.

In humans, the mortality of heat stroke can exceed 50% and there are no specific pharmacological treatments for heat stroke the most common therapeutic action being cooling [3]. The heat stress may result in cardiovascular disease such as heart failure, cardiac arrest and arrythmias [1], which accounts for majority of heat wave mortality [2]. It is likely that some of this mortality and disease may be due to increased vulnerability of heat stressed heart towards cardiovascular side effects of pharmaceuticals and other chemicals. Our data indicates that at least paclitaxel, lapatinib and phenanthrene had increased cardiac toxicity in zebrafish embryos under heat stress. This data implies that cardiotoxicity may be potentiated by heat stress.

Some medications such as psychotropic drugs may increase the patient′s susceptibility to heat stroke [22]. In humans, the mental disorders worsen with rising temperatures [5] and as extreme outcome the suicide risk is increased [23]. Human behavioural responses to heat stress are very complex and may involve many confounding factors, but at least in our zebrafish embryo experiments the heat stress itself affects behaviour. Aging is also a key risk factor for heat stroke [5] and rising global temperatures together with aging populations may result in increased prevalence of heat stroke. This indicates a burning need for new therapies for heat stroke. Although there is no approved pharmaceutical treatment for heat stroke, new potential drug targets are being explored [24]. The use of efficient screening models such as zebrafish embryo model could facilitate the discovery of such compounds. Zebrafish embryos are used actively as a tool to understand cardiovascular biology and disease [25,26], although physiological and anatomical differences between humans and zebrafish naturally exist. Differences in body temperature regulation (endothermic humans vs ectothermic zebrafish) is obvious. During heat stroke, however, the human body temperature regulation is disrupted. This actually then renders also humans in to an ectothermic state. The most effective current treatments to severe heat stroke – body cooling with cold water or ice packs [3] - is actually an ectothermic regulation of human body temperature.

Due to severity and increasing prevalence of heat stroke, the discovery of new pharmaceutical treatments for heat stroke could have large impact on human health. We envision that utilization of heat stress responses in zebrafish embryo could be utilized in discovery and investigation of new pharmacological treatments for heat stroke.

## Acknowledgements

We thank Zebrafish Core and Cell Imaging and Cytometry Core (both supported by Biocenter Finland) for technical assistance, services and infrastructure. We thank Ijlal Haider for assistance and Tuomas Kiviniemi for comments.

## Author contributions

Conceptualization; All authors, Formal analysis: KV, MA; Funding acquisition: IP; Investigation: All authors; Resources: IP; Supervision: IP; Visualization: KV, IP; Roles/Writing - original draft; all authors and Writing - review & editing: all authors.

